# There is no silver bullet: tiger corridors do not ensure multispecies carnivore connectivity

**DOI:** 10.64898/2026.07.05.736645

**Authors:** Divyashree Rana, Uma Ramakrishnan

## Abstract

Connectivity is critical to sustaining endangered carnivores in speciose yet fragmented landscapes such as in the global south. Corridors to mitigate fragmentation are designed based on charismatic species or habitat-based approaches, but their multispecies effectiveness for maintaining functional connectivity remains poorly tested. We combined landscape genetic analyses across five sympatric carnivores - *Panthera tigris, Panthera pardus, Prionailurus viverrinus, Felis chaus*, and Melursus *ursinus*, to evaluate how landscape features shape functional connectivity in a globally important felid landscape. We then assessed the efficiency of existing tiger corridors and single-species surrogates for maintaining multispecies functional connectivity. Species exhibited contrasting responses to landscape variables, producing distinct resistance surfaces and connectivity corridors. Spatial similarity of connectivity between species pairs was highly variable (r = 0.14-0.93), but no single species effectively captured connectivity patterns of the broader carnivore community (maximum mean overlap of <0.7 across species). Moreover, genetically optimized corridors were over 70% more efficient in capturing connectivity compared to existing tiger corridors, demonstrating mismatches between structural and functional connectivity. Our results highlight limitations of surrogate-based corridor planning and demonstrate that integrating multispecies functional connectivity can substantially improve conservation planning in human-dominated landscapes.

## Introduction

Connectivity is a fundamental property of ecological systems that enables the persistence of species across landscapes. As habitat fragmentation accelerates globally, maintaining and restoring connectivity has become a central goal of conservation planning (Beger et al., 2022; Fontoura et al., 2022). International policy frameworks, including the United Nations Sustainable Development Goals and the post-2020 Global Biodiversity Framework of the Convention on Biological Diversity, explicitly emphasize the importance of “well-connected systems” for halting biodiversity loss. Despite its importance, connectivity remains a complex and multifaceted concept that can be interpreted across spatial and temporal scales, from the physical arrangement of habitat patches (structural connectivity) to realized dispersal and gene flow among populations (functional connectivity) (Taylor et al., 2006).

Physically, wildlife corridors allow connectivity to be incorporated into spatial conservation planning. Corridors are typically defined as habitat linkages that facilitate the movement of organisms between otherwise isolated habitat patches in human-modified landscapes (Beger et al., 2022). Practical connectivity conservation therefore requires defining corridors and their networks, and guidelines for their design are embedded within policy and land-use planning frameworks worldwide (Keeley et al., 2019). Empirical and theoretical research helps identify corridors, and review of experimental studies have demonstrated the potential of these natural corridors to aid movement across fragmented landscapes (Hilty et al., 2012; Gilbert-Norton et al., 2010). However, in practice, most corridor designs rely primarily on structural features of landscapes (such as the presence of contiguous or forested habitat) rather than functional connectivity measures derived from movement or genetic data. However, advances in landscape ecology and molecular population genetics have increasingly enabled the quantification of functional connectivity across heterogeneous landscapes (Correa Ayram et al., 2016). Consequently, identifying and maintaining pathways that facilitate genetic connectivity is being recognised as the cornerstone of spatial conservation planning (Riginos & Beger, 2022).

A single species approach to connectivity is not unique. Conservation strategies have historically focused on a limited number of species assumed to represent broader biodiversity (Caro, 2010). Influential approaches such as umbrella, flagship, keystone, and indicator species emerged from this species-centric paradigm and continue to guide conservation policy and management (Simberloff, 1998). However, growing empirical evidence suggests that protecting habitat or connectivity for a single focal species does not necessarily safeguard ecological requirements of co-occurring species (Sibarani et al., 2019; Dutta et al., 2023). In response, connectivity planning has increasingly begun to incorporate multispecies perspectives (Wood et al., 2022). At most, multispecies corridor assessments rely on estimates of potential connectivity derived from habitat suitability surfaces (Cushman & Landguth, 2012; DeMatteo et al., 2017; Penjor et al., 2024), and so do not directly capture realized dispersal or gene flow across landscapes. Consequently, the effectiveness of corridors designed for focal or surrogate species in maintaining functional connectivity across multispecies communities remains largely untested. Empirical evaluations based on genetic connectivity are therefore critical for assessing whether single-species connectivity corridors can reliably support broader biodiversity conservation (Beger et al., 2022).

The Bengal tiger (*Panthera tigris tigris*) is the flagship for wildlife conservation in India, which supports the largest remaining global population of the species. Since the 1970s, sustained conservation efforts have led to substantial recovery of tiger populations, making India a global model for large carnivore conservation (Jhala et al., 2021). Beyond its symbolic value, the tiger also anchors national conservation policy. Under the guidance of the National Tiger Conservation Authority (NTCA), landscape-level planning frameworks identify networks of protected areas (tiger reserves) and habitat linkages (tiger corridors) considered essential for maintaining population connectivity in India (Gopal et al., 2007). These corridors, delineated using tiger occupancy estimates, are legal entities embedded within management plans with power to influence land-use decisions and infrastructure development across tiger landscapes. As a result, tiger-centric planning effectively functions as a surrogate framework for broader biodiversity conservation across many regions of India. More broadly, such single-species centric management, such as for tigers in India, typifies powerful yet untested silver bullet approaches in growing, resource-limited economies. But do such silver bullet approaches work? Do corridors designed for tigers adequately maintain connectivity for other carnivore species? Here, we use multispecies genetic data from non-invasive samples collected across the fragmented Terai Arc Landscape to evaluate the silver bullet hypothesis. By estimating functional connectivity for five sympatric carnivore species and comparing their inferred corridors, we assess whether tiger corridors can reliably serve as a surrogate for multispecies connectivity conservation.

## Methods

### Study area and sample collection

Field surveys were conducted across protected areas and surrounding multi-use landscapes encompassing Dudhwa Tiger Reserve and Pilibhit Tiger Reserve in the Central Terai region of northern India, a key portion of the Terai Arc Landscape. Sign surveys during the winters of 2022 and 2024 yielded non-invasive faecal samples from five sympatric carnivores: the Bengal tiger (*Panthera tigris*, TG), Indian leopard (*Panthera pardus*, LP), Fishing cat (*Prionailurus viverrinus*, FC), Jungle cat (*Felis chaus*, JC), and Sloth bear (*Melursus ursinus*, SB). Samples were collected using sterile swabs and preserved in Longmire’s buffer for downstream DNA extraction. Species identity of all samples was confirmed using PCR-based molecular assays.

### Genetic data generation

Species-confirmed samples were genotyped using species-appropriate sequencing approaches. Felid samples (TG, LP, FC, JC)) were processed using the Feliplex multispecies amplicon sequencing panel, which targets >30 cross-amplifying microsatellite loci to generate comparable multilocus genotypes across felid species as demonstrated in Rana et al. (2024). Samples from SB were processed using double-digest restriction-site associated DNA sequencing (ddRAD-seq) to generate genome-wide single nucleotide polymorphism (SNP) data following Tyagi et al. (2024). The resulting multilocus datasets were used to identify unique individuals for subsequent population and landscape genetic analyses.

### Landscape genetic modelling

For each species, spatial patterns of genetic differentiation among sampled unique individuals were used to infer landscape resistance to gene flow. Pairwise genetic distances were related to environmental and anthropogenic landscape variables (anthropogenic - agricultural density and distance to settlements, and environmental - distance to water and enhanced vegetation index) using a landscape genetic framework. Because species perceive landscapes at different spatial scales, each variable was evaluated at four spatial scales - 1, 2, 5, and 10 km. For each species, univariate models were used to estimate the direction and magnitude of resistance for each variable and identify the most informative spatial scale. Scale-selected variables retained from univariate analyses were combined in multivariate mixed-effects models to generate species-specific composite resistance surfaces. Model selection based on Akaike’s Information Criterion (AIC) was used to identify the most parsimonious multiscale resistance surface for each species.

### Corridor identification and multispecies comparison

Optimized resistance surfaces were used to estimate current flow across landscapes using protected areas as source nodes, generating species-specific connectivity maps and pairwise effective resistance. To understand connectivity patterns spatially, species-specific current flow was quantified using cumulative current distributions to assess the degree of spatial concentration versus diffusion. To assess multispecies connectivity by single species corridors, we compared spatial similarity in current flow using Spearman correlation and quantified the extent to which connectivity of one species was captured by areas identified for another. Next, species-specific corridors were “optimized” using an area-based threshold to match the area of “existing” tiger corridors in the landscape. The efficiency of the genetic optimization of the corridor was assessed based on proportion of connectivity captured by existing tiger vs optimized species-specific corridors. Lastly, using optimized current flows, new species-specific corridors were identified as pixels with the highest current density (top 5% and 10% quantile), to highlight potential multispecies connectivity hotspots for conservation planning in the landscape.

## Results

Of the 1,300 faecal samples collected across the study landscape, 1,068 were successfully genotyped and assigned to the target species (Figure SM1). These comprised 582 tiger, 194 leopard, 56 fishing cat, 47 jungle cat, and 189 sloth bear samples. Felid samples yielded multilocus genotypes across ∼35 microsatellite loci, while ddRAD sequencing of sloth bear samples produced 964 high-quality SNPs retained after filtering. Species-assigned multilocus genotypes were then used to identify unique individuals by removing duplicate detections of the same genotype. This resulted in a final dataset of 238 individuals across five carnivore species (tiger TG = 66, leopard LP = 35, fishing cat FC = 37, jungle cat JC = 35, sloth bear SB = 66) used for subsequent landscape genetic analyses.

### Effect of landscape on driving connectivity

Landscape variables influenced genetic connectivity differently across species in direction, spatial scale as well as magnitude (Figure 1). Agricultural density showed the strongest resistance for TG and SB, with rapidly increasing resistance at higher agricultural intensities, whereas JC and LP showed the opposite trend, with little impact on FC. Another anthropogenic variable, distance to settlements produced similar resistance gradients, with resistance decreasing closer to settlements for LP, JC, and SB. TG and FC, however, exhibited the opposite patterns, with resistance decreasing with increasing distance from settlements. Both the anthropogenic variables showed optimal effect at a broad spatial scale (10 km) for LP, JC, and FC, while effecting TG and SB at finer spatial scales (1 km, 2km, and 5 km).

**Figure 1.**
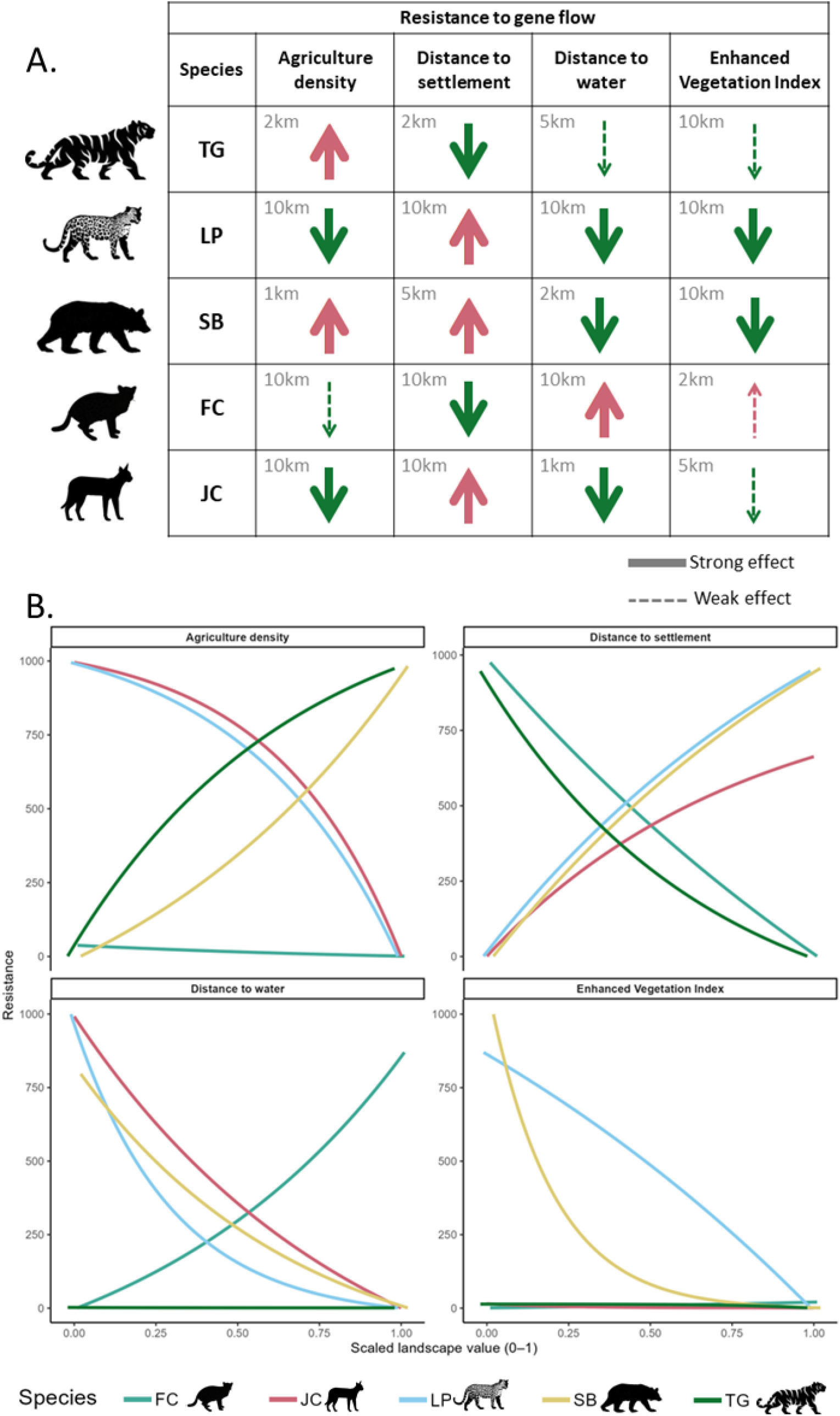
Comparative species-specific resistance patterns showing A. the optimized spatial scale and direction of effect and B. resistance curves for each landscape variable. In A, arrow direction indicates the effect on resistance to gene flow - red upward arrows represent increased resistance, whereas green downward arrows indicate reduced resistance. Solid arrows denote strong effects, while dashed arrows represent weak effects. In B, each coloured curve represents the optimized resistance relationship between a landscape variable and genetic connectivity for a focal species.

Natural variables showed comparatively less variation across species (Figure 1A). Distance to water strongly structured connectivity for most species at various scales, with resistance decreasing as distance from water increased, with the exception of fishing cats. EVI had relatively weak effects on connectivity for most species but showed stronger responses for SB and LP, with resistance declining rapidly with increasing vegetation cover for SB and more gradually for LP. Smaller optimal scales (2 km and 5 km) were observed for FC and JC, whereas larger optimal scales of 10 km characterized TG, LP, and SB with respect to EVI. Overall, resistance curves were most similar between JC and LP, and their connectivity along with FC responded to landscape variables at broader spatial scales. Together, these patterns highlight pronounced interspecific differences in the effect of landscape drivers shaping functional connectivity.

### Connectivity patterns and interspecies comparison

Species-specific resistance surfaces produced distinct connectivity patterns across the landscape. Connectivity distribution varied across species (Figure 2A). 8% of the landscape accounted for 50% of total cumulative connectivity for SB, compared to 13% for FC, 18% for TG, and approximately 27% for both LP and JC. These results indicate strong species-specific spatial differences in connectivity patterns, with SB showing the highest reliance on narrow connectivity pathways and LP and LC exhibiting more diffuse connectivity across the landscape.

**Figure 2.**
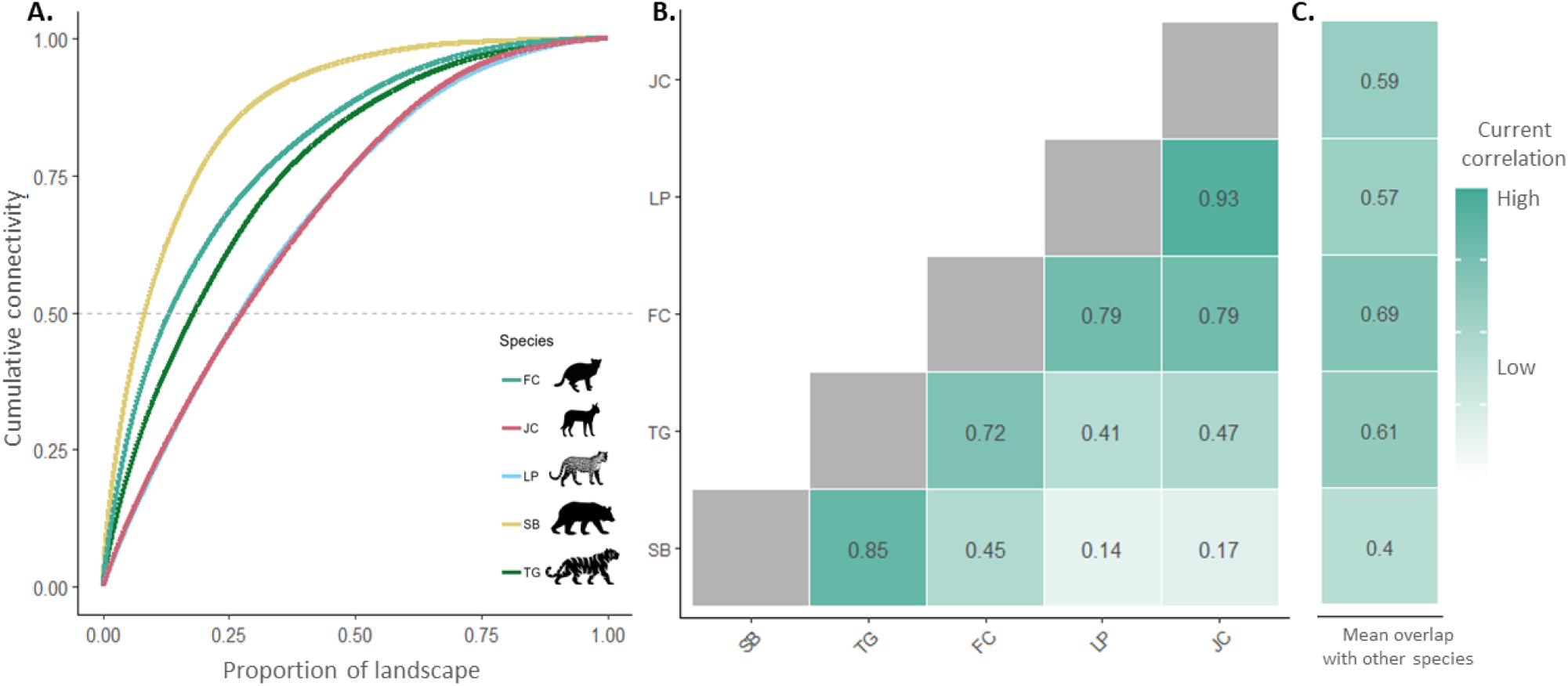
Species-specific connectivity patterns and their spatial similarity across the study landscape. (A) Connectivity distribution showing proportion of landscape accounting for percentage connectivity flow, (B) Heatmap showing pairwise correlation between species connectivity across the landscape, and (C) Mean current overlap of each species with others.

Correlation of current flows across the landscape for different species pairs ranged widely from 0.14 to 0.93 (Figure 2B). Despite differences in connectivity patterns across species, strong similarities were observed among certain species pairs. LP and JC showed the highest similarity in connectivity patterns (Spearman ρ = 0.93), followed by TG and SB (ρ = 0.85). In contrast, SB connectivity patterns showed least similarity with other species (least mean overlap of 0.4), specifically with LP (ρ = 0.14) and JC (ρ = 0.17). On average, FC showed intermediate similarity (0.45 - 0.79) with all species, with the highest mean overlap of almost 0.7.

### Efficiency of optimized corridors compared to existing tiger corridors for maintaining connectivity

The tiger corridors identified by NTCA capture an area of 549 km^2^ covering 2.92% of the entire study landscape. These existing tiger corridors capture less than 10% of the total connectivity for most species (TG = 8.3%, LP = 4.1%, JC = 4.4%, FC = 8%) with the exception of SB, capturing 13.8% of the total (Figure 3A). In comparison, the genetically optimized corridors of the same area cover a higher percentage of cumulative connectivity across all species, ranging from 7.3% to 26.3% (Figure 3B). Although optimized corridors were more efficient in capturing proportion of the connectivity than existing corridors, percentage gain varied across species. Proportionately, FC showed the maximum gain of over 135% with the least gain observed for TG at 69% (Figure 3B).

**Figure 3.**
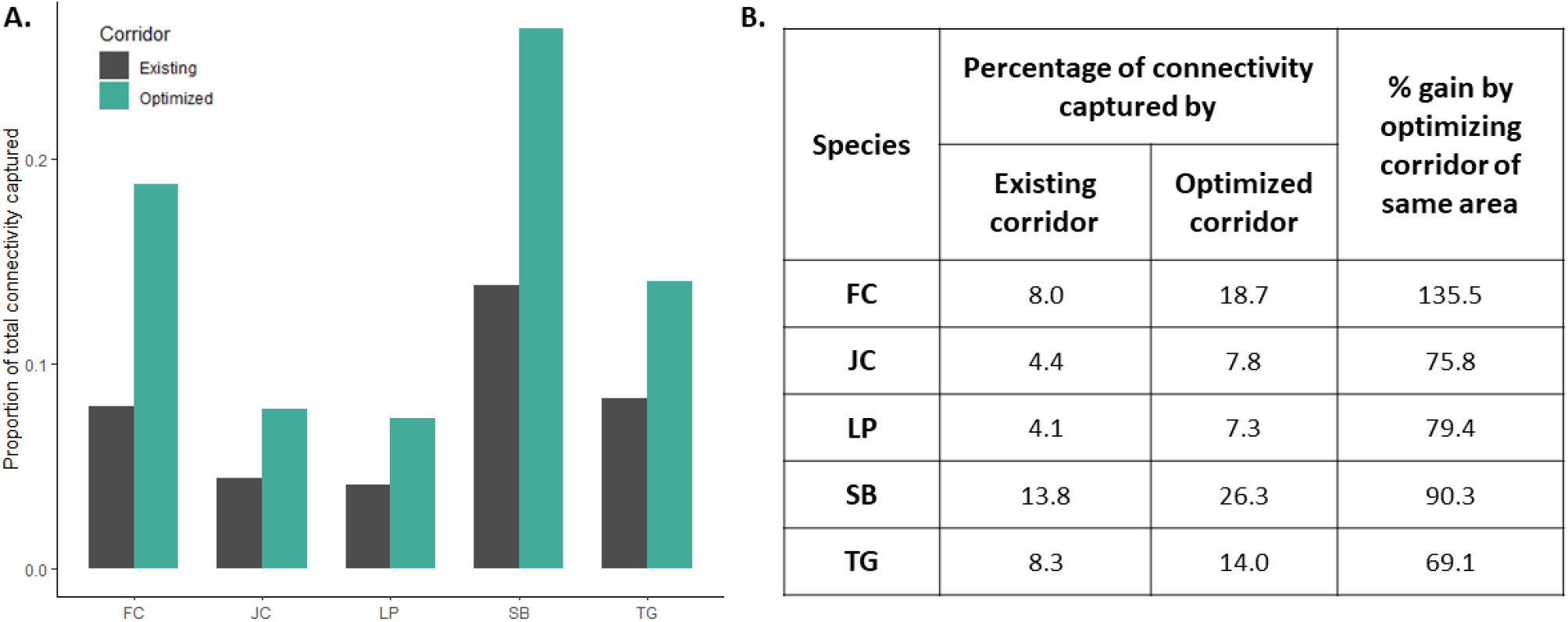
Connectivity captured by existing tiger corridors vs species-specific genetically optimized corridors. (A) Comparison of proportions and (B) Percentage captured and gained by optimization.

Genetically optimized connectivity corridors based on the current threshold of top 5% and 10%, although same in area covered, varied significantly across the species in their structure and location. Connectivity corridors for FC were the most contiguous (Figure 4A), with the low mean effective resistance across protected areas (1.49). These corridors primarily encompassed the surrounding multi-use landscapes including the meandering river basins. Corridors identified for LP and JC were spatially similar (Figure 4B-C), forming patchy connections between DDW and KAT and largely traversing agricultural multi-use areas, with least effective resistance values (<1). In contrast, TG corridors were more restricted, largely encompassing forested patches and wetland habitats, and exhibited the highest effective resistance values among species (mean = 6.91). Finally, SB corridors were the most distinct and spatially constrained, with no predicted corridor between DDW and KAT and a primary connectivity pathway following forested regions between PBT and KIS.

**Figure 4.**
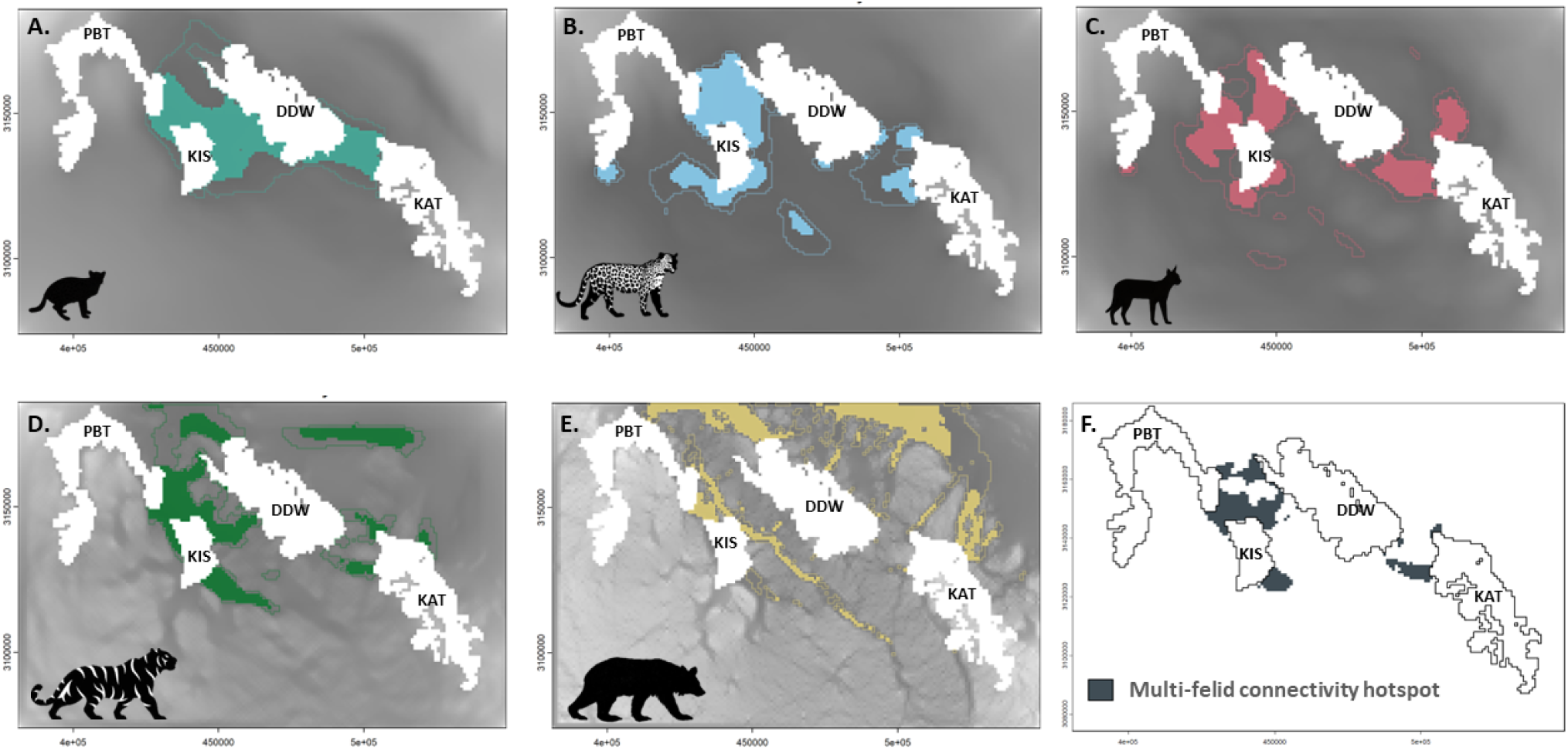
Species-specific critical corridors across the landscape derived from functional connectivity analyses. Panels show corridors (solid zone and outline showing top 5% and 10% connectivity hotspots) for (A) fishing cat FC, (B) leopard LP, (C) jungle cat JC, (D) tiger TG, (E) sloth bear SB, and (F) Critical wild felid connectivity hotspots.

Despite substantial differences in species-specific corridors, several areas emerged as zones of corridor convergence across multiple felid species (Figure 4F). These multispecies connectivity hotspots occurred primarily in forested regions linking major protected areas in the central portion of the landscape. However, large portions of species-specific corridors remained unique to individual species, indicating limited spatial overlap in functional connectivity requirements.

## Discussion

Maintaining ecological connectivity in increasingly human-modified landscapes is a central challenge for biodiversity conservation. Using a comparative landscape genetic approach across sympatric carnivore species, we evaluated how landscape features influence functional connectivity, and whether existing tiger corridors capture broader multispecies connectivity. We found substantial interspecific variation in landscape effects on gene flow, resulting in species-specific resistance surfaces and connectivity patterns. Consequently, overlap among current flow was moderate across species and no single species effectively represented connectivity patterns for the tested assemblage. Furthermore, the existing tiger corridors captured less than 15% of the cumulative connectivity across the landscape for all species including the tiger, and the genetically optimized corridors were over 70% more efficient. These findings highlight limitations of single-species and structurally derived corridor planning and emphasize the importance of multispecies evidence for connectivity conservation.

### Mismatch between structural and functional connectivity corridors

Most corridor identification approaches rely on habitat connectivity inferred from species presence or habitat suitability instead of functional pathways from movement (Brennan et al., 2020) or gene flow (Modi et al., 2025). For example, the existing corridor network primarily incorporates information on species presence, incorporating estimates of probability of occurrence to delineate potential connectivity pathways (Jhala et al., 2021). In contrast, our analyses explicitly modeled landscape resistance using patterns of genetic differentiation, providing an empirical estimate of functional connectivity.

The low efficiency of existing corridors in capturing connectivity in our results (Figure 3) suggests that corridors derived from habitat or occupancy information may not fully capture the pathways through which individuals actually move across complex landscapes. In our study system, optimized corridors frequently traversed multi-use areas, reflecting dispersal through heterogeneous landscapes. In contrast, existing tiger corridors were in fact more efficient for maintaining SB connectivity instead of the intended TG, as they were largely restricted to forested habitats. These patterns suggest that structurally derived corridors may preferentially capture habitat continuity while overlooking functionally important pathways via multiuse landscapes (Ghoddousi et al., 2021). This mismatch highlights an important scientific and policy gap, as many corridor networks incorporated into conservation planning rely primarily on habitat-based connectivity (Beger et al., 2022). We strongly advocate that functional connectivity data must be incorporated into corridor design, specifically in landscapes with fragmented natural habitats embedded in multiple land uses. Such evidence-based approaches will result in efficient corridor design and the desired connectivity outcomes (Merenlender et al., 2022).

### Limitations of single-species surrogates for multispecies connectivity

Using surrogate species to achieve multispecies conservation goals has long been a common strategy in conservation planning (Caro, 2010). However, growing evidence suggests that species often respond differently to landscape variables (Thatte et al., 2020; Modi et al., 2025), limiting the effectiveness of single-species approaches. As expected, our results demonstrate strong species-specific responses to landscape features (Figure 1), resulting in distinct connectivity corridors across the carnivore community.

In this landscape, agriculture dominated by sugarcane forms the primary matrix outside protected areas (Bista et al., 2024). Species for which agricultural density imposed strong resistance, such as TG and SB, exhibited high effective resistance and more restricted corridors between protected areas. In contrast, species known to tolerate human-modified environments, including LP and JC, showed lower resistance and broader connectivity through agricultural areas. These patterns are consistent with known species ecology (Chaudhary et al., 2025; Bandyopadhyay et al., 2026) and suggest that responses to anthropogenic land use may play a stronger role in shaping connectivity than traits such as body size alone.

Despite some similarities in resistance patterns among species, connectivity corridors remained highly species-specific. For example, although LP and JC exhibited comparable resistance responses, they only shared ∼50% overlap in critical corridor areas (Figure 4). Different frameworks have been proposed to identify surrogate species for multispecies connectivity (Meurant et al., 2018; Dutta et al., 2023). In our case, FC would emerge as a potential surrogate based on the highest mean current overlap with other species. However, optimized critical corridors show low congruence across species (Figure 4). These differences emerged even among closely related felids, highlighting the difficulty of representing multispecies connectivity with a single surrogate. Consistent with previous evaluations of umbrella species, surrogate species may provide a pragmatic starting point for conservation planning but rarely represent multispecies connectivity. (Wang et al., 2018; Sibarani et al., 2019; Wood et al., 2022).

### Future prospects for multispecies connectivity research and caveats

Comparative studies estimating functional connectivity across multiple species remain rare, largely due to the substantial data requirements of multispecies genetic datasets. Genetic research based on non-invasive samples is inherently challenging, and generating comparable genotypes across multiple species often relies on amplicon or reduced-representation sequencing approaches (Andrews et al., 2018). In this study we used a multispecies marker panel to generate comparable genetic data across felids, although such tools remain relatively novel and limited in their application (Rana et al., 2024). Continued development of similar multispecies genetic approaches that enable analyses at population and individual scales will be crucial for expanding comparative landscape genetic studies. Spatial prioritization is an important tool for identifying locations where conservation actions can benefit multiple species simultaneously (Wilson et al., 2009). With advances in genetic sequencing and connectivity modelling, landscape genetics provides a framework for evidence-based spatial planning, yet remains underutilized in conservation practice (LaPoint et al., 2015).

In contrast, analytical methods for resistance optimization have advanced rapidly in the past decade to infer realistic connectivity patterns (Liczner et al., 2024). However, most of these frameworks remain limited to single-species analyses, with multispecies patterns typically inferred through post hoc comparisons (Wood et al., 2022). Such approaches may also overlook the role of ecological interactions that shape species responses to landscapes. For example, in our study LP showed lower resistance through agricultural areas and near settlements (Figure 1), patterns consistent with the species’ well-documented tolerance of human-modified environments across India (Chaudhary et al., 2025; Kshettry et al., 2017). However, these patterns may also partly reflect spatial avoidance of habitats strongly used by tigers (Harihar et al., 2011), suggesting that realized connectivity may arise from a combination of landscape responses and interspecific interactions (Berger-Tal & Saltz, 2019). Hence, developing methods that explicitly integrate multiple species (and potentially their ecological interactions) within resistance optimization frameworks will be essential for designing connectivity strategies that more realistically reflect the complexity of ecological communities and improve the effectiveness of biodiversity conservation planning (Nielsen et al., 2017; Unnithan Kumar et al., 2022; Wood et al., 2022).

## Conclusions

India supports the highest diversity of wild felids globally (Dickman et al., 2015), and our study landscape represents an important regional hotspot for felid conservation. While conservation efforts in this region have largely focused on tigers, the landscape supports six sympatric felid species, four of which were included in our analysis. By integrating landscape genetic analyses across multiple species, our comparative framework not only identified species-specific resistance patterns but also revealed connectivity hotspots shared among several felids. Such a multispecies perspective provides a powerful pathway for translating species-specific insights into evidence-based fine-scale conservation strategies.

Unlike single-species approaches, multispecies analyses can help prioritize areas that maintain connectivity for entire ecological communities rather than individual focal species. Our findings highlight limitations of relying on single-species structural connectivity frameworks in diverse ecological communities. While tiger corridors have played a crucial role in guiding conservation investments for this apex predator across India, our results suggest that corridors optimized for a single species may not fully represent the connectivity requirements of sympatric carnivores. As conservation planning increasingly seeks to balance biodiversity persistence within human-dominated landscapes, integrating multispecies evidence will be essential to ensure that corridor networks sustain connectivity for entire ecological communities rather than individual flagship species.

## Acknowledgements

DR was supported by the TIFR-NCBS graduate program. We acknowledge the Panthera Corporation and International Bear Association for their funding support towards the project. Our gratitude extends to the National Centre for Biological Sciences for their institutional support as facilitated by UR, specifically the Next-Generation Genomics Facility (NGGF, Bangalore Life Science Cluster, BLiSC) for their assistance in data generation. The NCBS data cluster, supported under project no. 12-R&D-TFR-5.04-0900 by the Department of Atomic Energy, Government of India, was indispensable. We would like to thank the Uttar Pradesh Forest Department for providing necessary permits to collect non-invasive samples (permit no. 895 & 1148/23-2-12) and their on-site assistance during the field season. DR would like to thank the field team - student interns and local field assistants, who tirelessly contributed to the intensive field surveys. We would like to thank Dr. Samuel Cushman and Dr. Eduardo Eizirik for their suggestions in framing the project.

## Author Contributions

DR led project conceptualization, data collection, data generation, analyses, and manuscript writing. UR assisted in conceptualization while providing critical inputs and logistical support for the project. Both authors critically contributed to the drafts and approved the final version for publication. The authors declare no conflicting interests.

## Data sharing and accessibility

The data and codes are hosted in the primary author’s public GitHub repository and will be shared separately for the peer review process.

## Notes

https://github.com/divyashreerana/publications/tree/main/Rana%20et%20al.%202026_Tiger_corridors

### Competing Interest Statement

The authors have declared no competing interest.

