## Supplementary Material for "There is no silver bullet: tiger corridors do not ensure multispecies carnivore connectivity"

### SM1. Study area and sample collection

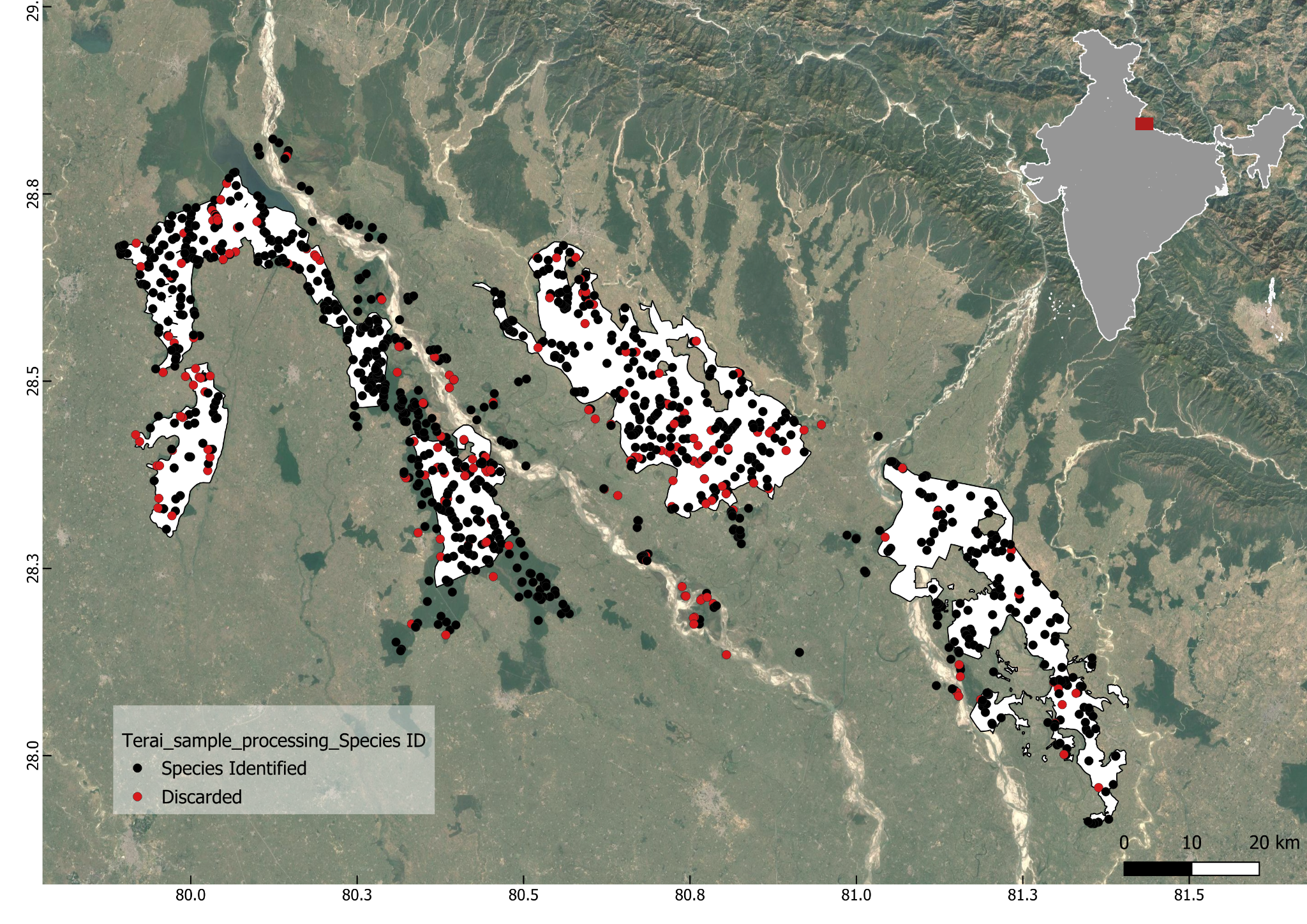

**Figure SM1. Sampling map marking genetically identified and discarded samples.**

*The red box within the India map on the top right corner marks the location of the study landscape within the country.*

### SM2. Genetic data generation

DNA was extracted using Qiagen Blood and Tissue Extraction kit following the manufacturers protocol and concentrations were measured using Qubit Fluorometer 4.0 (Invitrogen). The extracted DNA was first screened with a vertebrate 16S rRNA marker (Mukherjee et al., 2019). The amplified products were sequenced in the in-house Sanger sequencing facility and sequences were matched to the NCBI genetic repository for robust species identification.

#### SM2.1. Laboratory protocol for individual-level genetic data generation

Species-confirmed samples were genotyped using species-appropriate sequencing approaches. Felid samples were processed using the Feliplex multispecies amplicon sequencing panel (as explained below and in more details in Rana et al., 2024), which targets >30 cross-amplifying microsatellite loci to generate comparable multilocus genotypes across felid species.

The Feliplex markers were optimised for a two-step GT-seq library preparation protocol (Campbell et al., 2015) that includes an initial multiplexing reaction followed by an indexing reaction to attach unique barcodes to each sample. The PCR conditions for the first multiplexing reaction followed the protocol by De Barba et al. (2017) using QIAGEN Multiplex PCR Plus on extracted DNA, while the second indexing reaction followed the protocol by Natesh et al. (2019).

The primer pairs, modified by adding platform-compatible adapter regions, were divided into two genus-optimized pools of around 15-20 primer pairs each based on their amplification efficiency or read counts during optimization. The first multiplexing PCR was conducted separately for both pools with each sample, and the products were combined for the second indexing PCR. The indexed multiplexed samples were ultimately pooled in an equivolume manner and processed for 150bp paired-end sequencing on the Illumina NovaSeq6000 system. Each sample was sequenced three to five times using independent PCR replicates to generate consensus genotypes, as recommended for non-invasive samples (Taberlet et al., 1996).

Samples from sloth bears were processed using double-digest restriction-site associated DNA sequencing (ddRAD-seq) to generate genome-wide single nucleotide polymorphism (SNP) data. Owing to the presence of prey and bacterial contamination, faecal samples have low host DNA concentration. Thus, the samples genetically identified as belonging to sloth bears were enriched using methyl-CpG-binding domain (MBD) proteins to maximise host DNA concentration (Chiou & Bergey, 2016). The methylated bead-based enrichment method works on the biological principle that vertebrate DNA is more heavily methylated compared to microbial DNA, hence the magnetic beads coated with MBD2-Fc protein preferentially binds to methylation-rich vertebrate DNA (Wood & Zhou, 2016). For DNA enrichment, NEBNext Microbiome DNA enrichment kits were used (Feehery et al., 2013), for which MBD2-Fc magnetic beads were prepared by the conjugation of MBD2Fc protein and protein A magnetic beads. Prior to enrichment, the extract volume was reduced from 200 to 40uL using the CentriVap DNA concentrator (Labconco) to concentrate the DNA.

Post enrichment, the extracts were processed using double digested library preparation protocol (Peterson et al., 2012), while incorporating some modifications specific to low host DNA concentration samples (Tyagi et al., 2024). A combination of Sph1 and MluC1 restriction enzymes was used for digestion at 37°C for 3 hours followed by a 20 minutes heat-kill at 65°C, followed by adaptor ligation using T4 DNA Ligase at 23°C for 2 hours and 10 minutes at 65°C. Ligated products were purified to remove <100 bp fragments , followed by a 16-cycle indexing PCR (98°C for 30s, 65°C for 30s, 72°C for 30s and 72°C for 5min). Post indexing, fragments ranging from 200-500 bp were retained during purification (0.5X and 1X) with in-house prepared carboxyl group-coated SPRI beads. Individual sample libraries were quantified using a Qubit Fluorometer and their fragment size distribution was evaluated using the Agilent Tapestation.

DNA concentrations were measured after each processing step - i. extraction, ii. enrichment, and iii. library preparation - and products with unquantifiable DNA concentration were discarded. The remaining sample libraries were pooled in an equimolar fashion to generate comparable reads across varying sample concentrations. The pooled library was ultimately purified to retain fragments ranging from 200 to 500 bp, and was sequenced on the NovaSeq platform. Samples that yielded poor-quality data were re-sequenced and raw data for the same sample was merged to improve coverage.

The resulting sequencing datasets were used to identify unique individuals for subsequent population and landscape genetic analyses.

#### SM2.2. Bioinformatic processing for individual-level genetic data generation

The raw sequencing reads were trimmed to remove adapters and low-quality sequences (Q>30) using Trimmomatic (Bolger et al., 2014). The R1 and R2 reads were then aligned to generate consensus reads, which were demultiplexed for each primer pair using the *ngsfilter* function in OBITools (Boyer et al., 2016). A read count file was generated from the demultiplexed reads for each marker, containing the sample name, unique amplicon sequences, and their respective counts (read depth). Non-target reads without microsatellite repeats for the locus were excluded from further analysis.

Next, the number of microsatellite repeats in each sequence was counted, creating a table of unique sequences, their frequencies (read counts), and their microsatellite repeat numbers for each sample and marker combination. Only markers with a depth of >100 reads per sample were retained for further analyses. Putative allele sequences were identified based on the most abundant sequences containing the microsatellite motif, if it accounted for over 25% of the total reads. If a single allele sequence was identified, the PCR product was classified as homozygous, and as heterozygous if two such allele sequences were identified. PCR products with more than two putative alleles were deemed inconclusive. Alleles were identified across all loci for each PCR based on repeat sequence frequencies, resulting in the "genotype dataframe." A consensus genotype at each locus was called when at least three independent replicates produced matching genotypes, following De Barba et al. (2017). Genotyping error rates for allelic dropout and false alleles were calculated non-invasive samples by comparing alleles in each PCR replicate to consensus genotypes, following Broquet & Petit (2004).

Poor-quality samples and markers, defined as having over 40% missing data across samples, were discarded. Unique individuals were identified across all samples from nine felid species using GenAlEx 6.5 (Peakall and Smouse, 2006), based on matching probabilities of multilocus consensus genotypes (Waits et al., 2001). Samples with three or fewer locus mismatches, ignoring missing data, were considered recaptures of the same individual. The frequency of null alleles was estimated across loci for all species using the Brookfield (1996) method, implemented through the PopGenReport R package (Adamack & Gruber, 2014). Genus-specific genetic clustering of identified individuals was examined using Principal Component Analysis (PCA) and Discriminant Analysis of Principal Components (DAPC) via the adegenet R package (Jombart, 2008). The variation explained by the first two principal component axes was used to visualise genetic differentiation between individuals from different species within each genus.

For ddRAD sequences from sloth bear samples, demultiplexed reads were quality checked using *FastQC* and compiled to a single .html report using *MultiQC* (Ewels et al., 2016). Adaptor sequences, low quality bases and very short reads were removed using *Trimmomatic*. As the reads were generated from faecal samples, contamination with prey as well as bacterial DNA was a major concern despite the enrichment. Hence, the raw reads were trimmed for adapters, using Trimmomatic, and then mapped to a reference genome of the closest related species - sun bear Helarctos malayanus (GCA_028533245.1) (divergence time of ~3.5 million years with the last common ancestor) (Kumar et al., 2017), in absence of a sloth bear reference genome, using *BWA-MEM* (Li, 2013). The bam files, retaining only the mapped reads, were then converted to a .fastq file using *BEDTools* (Quinlan et al., 2010) to be used for downstream processing.

The filtered reads were then processed using the *ipyrad* bioinformatic pipeline with 28 default parameters to conduct de-novo local assembly and analyze the ddRAD sequence data (Eaton & Overcast, 2020) using default parameters to generate a Variant Call Format (VCF) file with SNPs. The filters ensured a minimum depth of six bases, 85% clustering threshold, and retention of loci present in a minimum of four samples. Additionally, reads with more than five low quality bases, *phred* score of less than 33, or length less than 35 were removed, and adaptors were trimmed. Further details of ipyrad parameters used can be found in the Table SM1.

**Table SM1: Ipyrad parameter list with values and paths provided**

| **PARAMETER NAME** | **PARAMETER**  **DESCRIPTION** | **VALUES AND PATHS PROVIDED** |
| --- | --- | --- |
| assembly_name | Used to assign a prefix to all output files | sbanalysis |
| project_dir | Path to a directory where files are to be stored | slothbearanalysis |
| raw_fastq_path | Path to non-demultiplexed files | - |
| barcodes_path | Path to file containing barcodes | - |
| sorted_fastq_path | Location to demultiplexed files | /home/uramakri/utkarshschauhan/Judi/sbanalysis_filtered/inputsb/*.fastq.gz |
| assembly_method | Offers choice from either denovo clustering or reference mapping | denovo |
| reference_sequence | path to where the reference sequence is downloaded | - |
| datatype | Choice from 6 different datatypes used in RAD-seq | pairddrad |
| restriction_overhang | Detect and filter out adaptor sequences | CATGC,AATT |
| max_low_quality_bases | Sets the upper limit for the number of ambiguous sites (N) allowed in reads | 5 (allows upto 5 N) |
| phred_score_offset | Reads filtered and trimmed if quality scores are not a certain value | 33 (default) |
| mindepth_statistical | Minimum depth of statistical base calling | 6 (default) |
| mindepth_majrule | Minimum depth of majority rule base calling | 6 (default) |
| maxdepth | Sets the maximum value beyond which clusters would be excluded | 10000 (default) |
| clust_threshold | Similarity levels of two sequences being clustered together. | 0.85 (default) |
| max_barcodes_mismatch | Number of mismatches allowed | 0 |
| filter_adaptors | Levels of strictness in filtering out illumina adaptors | 2 (strict filtering) |
| filter_min_trim_len | Minimum length of reads after trimming to be present | 35 (default) |
| max_alleles_consensus | Highest number of unique alleles present depending on ploidy of individual | 2 (since diploid) |
| max_Ns_consens | Maximum percentage of uncalled bases allowed in a consensus | 0.05 (default ) |
| max_Hs_consens | Maximum number of heterozygous bases allowed in consensus | 0.05 (default) |
| min_samples_locus | Lower threshold for number of samples containing data at a particular locus | 4 (default) |
| max_SNPs_locus | Maximum SNPs allowed in final locus | 0.2 (default) |
| max_Indels_locus | Maximum insertions and deletions allowed in final locus | 8 (default) |
| max_shared_Hs_locus | Maximum number of shared polymorphic sites in a locus | 0.5 (default) |
| trim_reads | Specific trimming of N bases at the beginning or end of a sequence | 0, 0, 0, 0 (no extra trimming required) |
| trim_loci | Specific locus edge trimming | 0, 0, 0, 0 (no extra trimming required |
| output_formats | Choose from various output formats that can be user-defined based on type of analysis | v (can specify up to 3) |
| pop_assign_file | Assigns unique identify to samples | - |
| reference_as_filter | Used to remove sequences mapped to a reference genome | - |

Reduced-representation sequencing approaches such as ddRAD may experience allele or locus dropout due to restriction-site polymorphisms or uneven sequencing coverage. To minimize such effects, the output VCF file containing the variants was further filtered using *vcftools*. The final filtered vcf contained only biallelic sites with a minimum depth of six and maximum depth of 1000 copies and minor allele count (MAC) of more than three to avoid very rare or technical variants. Lastly, samples with over 60% and loci with over 40% missing data were dropped to remove poor quality data and reduce overall missingness.

### Landscape genetic modelling

For each species, spatial patterns of genetic differentiation among sampled individuals were used to infer landscape resistance to gene flow. Pairwise genetic distances were related to environmental and anthropogenic landscape variables using a landscape genetic framework. We evaluated two anthropogenic variables (agricultural density and distance to settlements) and two environmental variables (distance to water and enhanced vegetation index). Because species may perceive landscapes at different spatial scales, each variable was evaluated at four spatial scales (1, 2, 5, and 10 km). Univariate models were used to estimate the direction and magnitude of resistance for each variable and identify the most informative spatial scale. Variables retained from univariate analyses were combined in multivariate mixed-effects models to generate species-specific composite resistance surfaces. Model selection based on Akaike’s Information Criterion (AIC) was used to identify the most parsimonious multiscale resistance surface for each species.

To understand the influence of landscape features on observed spatial population structure for the species, a landscape genetic approach was implemented. This involves parametrization of resistance surface for each landscape feature determining the resistance offered by the latter to movement of the species using pairwise genetic distances. Based on sloth bear ecology, landscape properties, and published literature, four uncorrelated landscape variables (<|0.6|) (distance to water, agriculture land cover density, enhanced vegetation index, and distance to human settlement) relevant to understanding the connectivity of sloth bear populations were selected (Table SM2) (Dutta et al., 2015; Thatte et al., 2019; Malik et al., 2023). All variables were reclassified and recalculated as explained in Table 4.1. Distance layers were generated using the “Proximity (Raster Distance)” function in QGIS with a binary target layer as the input raster. As studies have shown that species-landscape interactions are scale-dependent (Cushman & Landguth, 2010; Suárez-Castro et al., 2018), all variables were processed at four spatial scales (1 km, 2 km, 5 km, and 10 km) with a moving window focal mean using the terra package in R (Hijmas et al., 2022).

**Table SM2. Description and details of landscape variables used in the study.**

| **Variable** | **Description** | **Source** | **Details** |
| --- | --- | --- | --- |
| Distance to water | Distance from the centre of each pixel to the center of the nearest permanent inland water | Google Earth Engine  (ee.ImageCollection("GLCF/GLS_WATER")) | Binary water layer based on category 2: “water” |
| Agriculture landcover density | Percentage of area occupied by agriculture landuse | Google Earth Engine  (ee.ImageCollection("ESA/WorldCover/v100")) | Per pixel density for category 40: “agriculture” |
| Enhanced vegetation index (EVI) | Satellite-derived metric for vegetation density correcting for atmospheric noise | Google Earth Engine  (ee.ImageCollection("MODIS/MOD09GA_006_EVI") | Raster scaled by a factor 0.0001 |
| Distance to human settlement | Distance from the centre of each pixel to the center of the nearest to human settlement | Google Earth Engine  (ee.ImageCollection("JRC/GHSL/P2023A/GHS_BUILT_C")) | Binary settlement layer generated by combining categories 12-25 (builtup area) |

A multi-scale multi-surface optimization approach was used to understand the effect of landscape features in governing gene flow, and to create a composite resistance surface using resistanceGA package in R (Peterman, 2018). This was based on a mixed effect linear mixed effects model fitted to the genetic data with a maximum likelihood population effects parameterization (MLPE), which has been identified as the optimal choice for landscape genetic analysis (Shirk et al., 2018).

As our target carnivores are primarily solitary in nature, individual-based analysis was implemented to understand landscape effects on gene flow across sampled individuals. Genetic distances were calculated between unique individuals using the “*diss.dist*” function of the *poppr* package in R. The rasters for the landscape variables were converted to ascii format using the terra package in R. With genetic data as the response and effective cost distance within each landscape variable as predictors, single surface models were optimized with default parameters to attain lowest AIC values within a maximum of 1000 iterations. Hence, at every scale, each landscape variable was optimized to identify the transformation type, shape of the transformation, and the maximum resistance offered using the “SS_optim” function in the ResistanceGA package in R. Therefore, using univariate optimization, scale and best-suited model parameters for each landscape variable were identified.

The optimized single surface parameters were used to understand the influence of landscape variables on spatial genetic population structure. These optimized parameters for each scale-selected raster variable were further used to generate composite surfaces with all combinations of the five variables to optimize the composite landscape resistance layer using the “MS_optim” function in the ResistanceGA package in R. The optimized multisurface raster layer with least AIC value was used to model functional connectivity across the landscape. Using the optimized multisurface resistance layer and protected area node raster, the connectivity was modelled for sloth bears using the pairwise modelling mode with default parameters in *Circuitscape 4.0* software (McRae et al., 2009).

### Corridor identification and multispecies comparison

Tiger corridors downloaded as a KML from NTCA official website (<https://ntca.gov.in/dss/#decision-support-system>). The downloaded file was clipped to the study area extent to compare existing tiger corridors to our optimized species-specific corridors.
